# EPIP: MHC-I epitope prediction integrating mass spectrometry derived motifs and tissue-specific expression profiles

**DOI:** 10.1101/567081

**Authors:** Weipeng Hu, Si Qiu, Youping Li, Xinxin Lin, Le Zhang, Haitao Xiang, Xing Han, Sitao Zhu, Lei Chen, Sha Li, Wenhui Li, Zhe Ren, GuiXue Hou, Zhilong Lin, Jianliang Lu, Geng Liu, Bo Li, Leo J Lee

## Abstract

**Background:** Accurate prediction of epitopes presented by human leukocyte antigen (HLA) is crucial for personalized cancer immunotherapies targeting T cell epitopes. Mass spectrometry (MS) profiling of eluted HLA ligands, which provides high-throughput measurements of HLA associated peptides *in vivo*, can be used to faithfully model the presentation of epitopes on the cell surface. In addition, gene expression profiles measured by RNA-seq data in a specific cell/tissue type can significantly improve the performance of epitope presentation prediction. However, although large amount of high-quality MS data of HLA-bound peptides is being generated in recent years, few provide matching RNA-seq data, which makes incorporating gene expression into epitope prediction difficult.

**Methods:** We collected publicly available HLA peptidome and matching RNA-seq data of 34 cell lines derived from various sources. We built position score specific matrixes (PSSMs) for 21 HLA-I alleles based on these MS data, then used logistic regression (LR) to model the relationship among PSSM score, gene expression and peptide length to predict whether a peptide could be presented in each of the cell line. We further built a universal LR model, termed Epitope Presentation Integrated Prediction (EPIP), based on more than 180,000 unique HLA ligands collected from public sources and ~3,000 HLA ligands generated by ourselves, to predict epitope presentation for 66 common HLA-I alleles.

**Results:** When evaluating EPIP on large, independent HLA eluted ligand datasets, it performed substantially better than other popular methods, including MixMHCpred (v2.0), NetMHCpan (v4.0), and MHCflurry (v1.2.2), with an average 0.1% positive predictive value (PPV) of 52.01%, compared to 37.24%, 36.96%, 24.90% and 23.76% achieved by MixMHCpred, NetMHCpan-4.0 (EL), NetMHCpan-4.0 (BA) and MHCflurry, respectively. It is also comparable to EDGE, a recent deep learning-based model that is not publicly available, on predicting epitope presentation and selecting immunogenic cancer neoantigens. However, the simplicity and flexibility of EPIP makes it easier to be applied in diverse situations, and we demonstrated this by generating MS data for the HCC4006 cell line and adding the support of HLA-A*33:03 to EPIP. EPIP is publicly available as a web tool < http://epip.genomics.cn/>.

**Conclusions:** we have developed an easy to use, publicly available epitope prediction tool, EPIP, that incorporates information from both MS and RNA-seq data, and demonstrated its superior performance over existing public methods.

## Background

T cell epitopes, which are short peptides presented by major histocompatibility complex (MHC) molecules on the cell surface and recognized by T-cell receptors (TCRs), lie at the heart of the human immune system that can remove infected and malignant cells. With the rapid progress of cancer immunotherapies, the mechanistic understanding, computational prediction and clinical manipulation of T cell epitopes have gathered renewed and widespread interests [1–4]. It is well known that peptide binding to MHC molecules is the most selective step in the antigen presentation pathway [5], and intensive computational efforts have been exerted to predict this binding process based on peptide sequences (allele-specific models) or peptide and the corresponding human leukocyte antigen (HLA) sequences (pan-specific models). Traditionally, these models are trained on data accumulated from various types of *in vitro* binding assays, which can provide quantitative binding affinity measurements between pre-selected peptides and HLA-I or HLA-II molecules. However, the requirement of synthesizing peptides beforehand limited the ability of these assays to perform unbiased, high-throughput screening of the vast peptide space, and it also did not account for the antigen-loading process *in vivo* [6]. The recent advance of mass spectrometry (MS) profiling of HLA ligands, mainly for HLA-I alleles at the moment, overcomes most of these limitations by providing *in vivo* measurements of peptides presented by MHC molecules on the cell surface in a high-throughput manner, although the resulting measurements are only qualitative (binary).

The value of MS data as contributing to the ultimate prediction of peptide immunogenicity has been increasingly recognized [6, 7], and as a result, a number of recently developed peptide-MHC binding prediction methods incorporated some form of MS data, in addition to the traditional binding assay data [5–8]. For example, NetMHCpan-4.0 trained an ensemble of two hidden layer neural networks on both affinity and MS data, and was able to make predictions on both types of outputs. MHCflurry adopted a deep learning framework containing both locally connected and fully connected layers and was mainly trained on affinity data, but used MS data at the model selection stage, while MS data may play an even bigger role in its future releases. Other methods chose to rely on MS data alone [7, 9, 10], such as MixMHCpred [9, 10], which built allele-specific position score specific matrixes (PSSMs) after deconvolving mixed-allele MS data, and the recent EDGE method [7], which used a comprehensive, deep learning model that pooled together large sets of mono-allele and mixed-allele MS data as well as matching RNA-seq data. With the increasing amount of MS data becoming available, it is our view that it might be better to build separate models for binding affinity (based on binding assay data) and epitope presentation (based on MS data) since the underlying data generation processes are different. These two related pieces of information could be further combined at a later stage to contribute to the prediction of immunogenicity. The presentation of a peptide by MHC not only depends on information contained within the protein sequences but is also modulated by other factors such as protein abundance, localization and turnover [11, 12]. Among these, the corresponding gene expression levels can be handily profiled by RNA-seq and have been shown to significantly improve prediction accuracy, first by the MS IntrinsicEC method on mono-allele cell lines [12], then more comprehensively by EDGE [7]. Unfortunately, both methods are not publicly available.

We set out to develop an effective, flexible and publicly available HLA-I epitope prediction method that takes advantage of large sets of MS and RNA-seq data. We first collected public MS and matching RNA-seq datasets for 21 HLA-I alleles, which frequently appear in the European or Asian population. By carefully examining the contributions of PSSM scores, gene expression levels and peptide length features to epitope presentation, we were able to effectively model these variables with a universal logistic regression (LR) model. We then trained such a LR model, termed Epitope Presentation Integrated Prediction (EPIP), to support 66 common HLA-I alleles. In the remainder of this article, we first describe our design rationale in detail, followed by comprehensive evaluation of EPIP on large, independent MS and immunogenic datasets as well as a user case showing its extendibility, before concluding with discussions on future improvements and research directions. We believe EPIP is a timely contribution to the research community of cancer immunotherapies and beyond.

## Methods

### Collection of MS and RNA-seq data

We collected matching MS and RNA-seq data from 16 mono-allele HLA-A and HLA-B cell lines [12] and 18 mixed-allele cell lines [13] to build the EPIP model. We also collected MS data of 15 mono-allele HLA-C cell lines from Di Marco et al [14], deconvolved MS data of 26 more HLA alleles from SysteMHC [15] and a small portion of mixed-allele MS data released with EDGE [7] to expand the allele coverage of EPIP. To perform independent evaluation of EPIP, we further downloaded MS data from Trolle et al [16] and Bassani-Sternberg et al [11], in addition to test data reserved from Pearson et al [13] and Bulik-Sullivan et al [7]. Detailed information and usage of all collected MS data are provided in Table S1.

### MS profiling of HCC4006 HLA-I peptidome

HLA-I peptidome samples for HCC4006 were prepared according to Bassani-Sternberg et al [11]. They were then analyzed by LC-MS/MS to obtain peptide MS spectra, and full details of the experimental procedure are provided in the Supplementary Material. We employed the MS-GF+ search engine version 2018.07.17 [17] to search the peak lists against the UniProt databases (161,521 entries for human as of December 2017) and a file containing 245 frequently observed contaminants such as human keratins, bovine serum proteins, and proteases. N-terminal acetylation (42.01 Da) and methionine oxidation (15.99 Da) were set as variable modifications. The enzyme specificity was set as unspecific. The initial allowed mass deviation of the precursor ion deviation was set to 10 ppm. Possible peptide lengths were restricted to 8 to 15. Possible precursor charges were restricted to 2 to 5. Range of allowed isotope peak errors was restricted to −1 to 2. Percolator version 3.02.0 was applied as a post-processing step for result filtering [18]. The *q* value cutoff was set to 0.01. From the “pout.tab” output file produced by percolator, hits to the contaminants were eliminated.

### Data and features of EPIP models

From the matching MS and RNA-seq data collected from public sources [12, 13], we first built separate LR models, EPIP_s, to predict epitope presentation for 21 alleles and tissues. This included all 16 mono-allele data profiled in Abelin et al [12], as well as five more alleles (HLA-A*11:01, HLA-A*32:01, HLA-B*15:01, HLA-B*40:01, HLA-B*07:02) selected from Pearson et al [13], which frequently appear in the Chinese population [19]. All RNA-seq datasets were downloaded and re-processed with the same pipeline to ensure consistency. Specifically, raw fastq files were cleaned by fastp (version 0.18.0) [20]and aligned to human genome assembly GRCh38 and its associated transcriptome (Ensembl Release 92) with STAR (version 2.5.3a) [21]; gene expression (transcripts per million or TPM) was quantified by RSEM (version 1.3.0) [22] based on the alignment.

EPIP_s uses three features to model epitope presentation: peptide motifs as modelled by PSSM, peptide lengths and the corresponding gene expression levels. For mono-allele MS data, we built PSSMs directly, as described in Liu et al [23]. For mixed-allele MS data, we first performed motif deconvolution by GibbsCluster [24] before building PSSMs. Peptides from deconvolved motifs that matched the HLA motifs of interest generated from SysteMHC data were retrieved. For each allele, length-specific PSSMs were built for those that have at least 100 peptides, up to the most abundant three lengths (9-11-mers or 8-10-mers, depending on the allele, details in Table S1), and this allowed us to make use of 74%-96% of MS profiled peptides across different alleles. To fully utilize the training data and avoid overfitting, we used the stacking method [25] to build PSSM (see supplementary materials, Fig S1). For each HLA allele, we also obtained an empirical probability mass function (pmf) to depict the length distribution of 8-15-mers based on the allele-specific MS data, and the peptide length feature is the corresponding probability in the empirical pmf of peptide length distribution. Expression values of a peptide was calculated by summing up the expression levels of all transcripts containing the peptide (in TPM) before transformed by log2(TPM+1). For the mono-allele cell lines in Abelin et al [12], transcript expression levels were taken as the average of the four cell lines with RNA-seq data available; similarly, for the mixed-allele cell lines in Pearson et al [13], transcript expression levels were averaged over the 10 cell lines with RNA-seq data available. All three features were further normalized to be zero mean, unit variance Gaussian before using as inputs to a LR model with weak L2 regularization implemented in scikit-learn 0.19.0 [26].

MS profiled peptides can only provide a positive training set for our model. To generate the negative set, we randomly sampled the human peptidome to obtain peptides of the matching length and remove those that overlap with the positive set. Adopting the same strategy as in NetMHCpan 4.0, we used more negative peptides than positive ones to train EPIP_s. For example, if the positive set contains 9-11-mers with the most abundant 9-mers having a size of *n*, we then generate a negative set with 10*n* peptides for each length (30*n* in total).

To build the full EPIP model for the above 21 alleles from different cell lines, we pooled all the PSSM scores, peptide length distributions and corresponding gene expression levels together to train a single LR model with the same training data as EPIP_s. Five-fold cross validation was used to evaluate EPIP during training and to compare EPIP and EPIP_s, while the full training set was used to build the final EPIP model and it was evaluated on independent test data. To expand the number of HLA alleles that EPIP can support, mono-allele HLA-C MS data from Di Marco et al [14], deconvolved MS data from SysteMHC [15] and some mixed-allele MS data released by EDGE [7] were used to learn PSSMs for these additional HLA alleles. For the mono-allele HLA-C MS data from Di Marco et al, we deconvolved the unfiltered list of peptides to remove the B35 and C04 motifs through GibbsCluster [24] before using them to train PSSMs. We further generated additional MS-data from the HCC4006 cell line so that EPIP can support HLA-A*33:03, a high frequency allele with 8.03% prevalence in the Chinese population [19].

### Immunogenicity evaluation

The immunogenic dataset used in this paper is the same as in Bulik-Sullivan et al. [7], which contains 2,023 assayed mutations from 17 patients with annotated HLA information from four previous studies [27–30]. In three of the four studies [27, 28, 30], mutations were tested using 25mer tandem minigene (TMG) assays. For each mutation in these datasets, we used EPIP, MixMHCpred 2.0, NetMHCpan4.0-EL, NetMHCpan4.0-BA, MHCflurry 1.2.2 to make predictions with respect to the HLA alleles of each patient for overlapping 8-11mers that harbor the mutation, and the score of the mutation is taken as the best score among these peptides (EPIP: highest presentation probability; MixMHCpred 2.0: highest predicted score; NetMHCpan4.0-EL and NetMHCpn4.0-BA: lowest rank score; MHCflurry 1.2.2: minimum binding affinity). We then ranked the mutations according to the best scores from each software. For the fourth study [29], we ranked mutations by taking the best scores from each software across all mutation-spanning peptides tested in the tetramer assays. To compare EPIP with EDGE on predicting immunogenic peptides, we excluded the peptides whose immunogenicity was undetermined as described in Bulik-Sullivan et al. We retrieved a total of 31 immunogenic epitopes from Tran et al., 216 Gros et al. and Stronen et al., which were not provided by Bulik-Sullivan et al and slightly different from the number 29 reported in Bulik-Sullivan et al^1^. Since the epitope prediction scores by EDGE was not available, we directly compared its immunogenic prediction results with EPIP.

## Results

### Construction and analysis of EPIP models

After collecting matching MS and RNA-seq data from public sources, we first developed separate LR models for each of the 21 tissues and alleles (termed EPIP_s) to predict epitope presentation by combining motif, peptide length and gene expression features. Full details of the datasets and features are described in Methods, and the overall process is sketched in Figure 1A. According to previous studies [12, 32–34], about one in a thousand peptides generated from the human proteome can be presented by MHC. Therefore, we further generated 999-fold decoy peptides that are not in the training set to evaluate our models. Similar to Abelin et al [12], the main evaluation criterion used throughout the paper is 0.1% positive predictive value (PPV), which measures the proportion of truly presented peptides in the top-scoring 0.1% ones found by our model. The 0.1% PPV is both a more realistic measure for immunopeptidome prediction and a more stringent one than many other popular metrics, such as ROC-AUC and F1 scores, for this highly biased classification task.

**Fig. 1.**
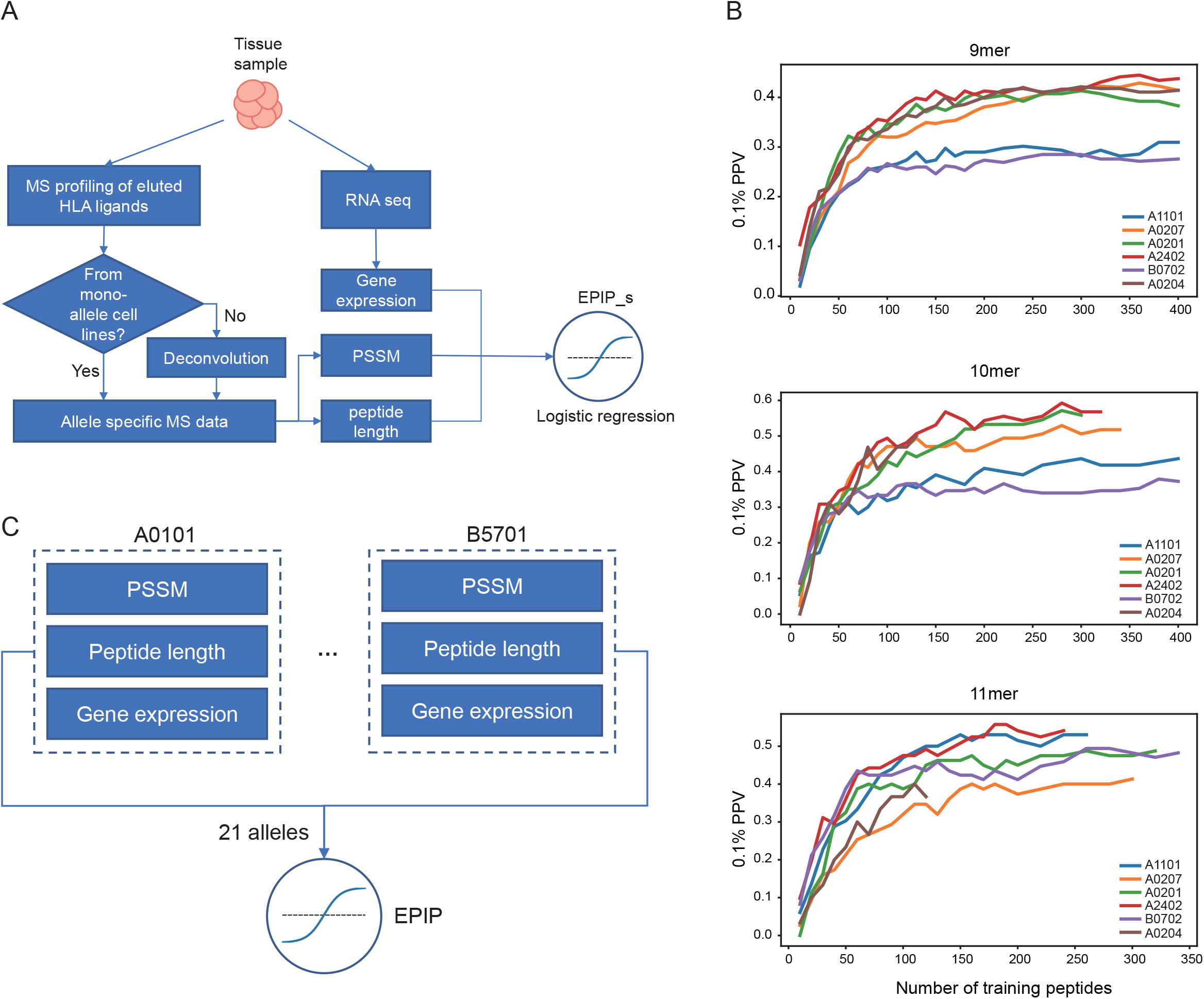
Construction of EPIP_s and EPIP. a) Construction of allele and tissue-specific EPIP (EPIP_s). EPIP_s was built for 21 alleles that have matching expression profile. b) Saturation analysis of PSSM models. The PSSM models were trained and tested on 9-11mer peptides independently for six HLA alleles; evaluated with 0.1% PPV, the performances of PSSM improved with increasing training data, and reached a plateau with about 100-200 peptides, depending on the specific alleles. c) Construction of EPIP. To build the full EPIP model, we pooled peptides from 21 alleles encoded by PSSM, expression and length features to fit a single LR model

We used PSSM, a simple model that can learn patterns in biological sequences, to model peptide motifs since it has previously been widely used in modelling MHC binding peptides [35–37], is easy to visualize, and seems to have adequate representational power. It is also a data efficient model due to its simplicity. To show this, we used subsampling to build PSSM models for different lengths of peptides to predict epitope presentation and tested their performances (0.1% PPV) on various alleles. As shown in Figure 1B, the performances of PSSM models reached saturation when having 100-200 peptides in the training set. Therefore, we selected 100 as a cutoff to build length-specific PSSM models for up to three most abundant lengths in each allele (9-11-mers or 8-10-mers, depending on the allele, details in Table S1), and the model would also be restricted to make predictions on these peptide lengths.

Although it is reasonable to assume that the relationship among motif, peptide length and gene expression is a general one across different tissues (cell lines) and alleles so that data from these different sources could be pooled together to train a general LR model (as outlined in Figure 1C), building EPIP_s first allowed us to rigorously analyze and quantify this. We first examined the distribution of optimized model parameters (weights and biases) among these 21 EPIP_s models. As shown in Figure 2A, the corresponding model parameters distributed quite tightly. Since the features have been normalized prior to model training, the relative magnitudes of model parameters also displayed the contribution of each feature, where PSSM played the most important role, followed by expression levels and peptide lengths (the average weights of PSSM, expression level and peptide lengths are 4.30, 1.37, 0.37, respectively). Furthermore, when displaying these 21 sets of weights and biases as hive plots (Figure 2B), they share very similar inter-relationships and seem to be mostly differ by a scaling factor, which is likely caused by somewhat different influences of L2 regularization in each model. This provided us with a strong motivation to build a universal LR model, EPIP, to account for the contribution of PSSM, peptide length and gene expression across these 21 tissues and alleles, to compare against the EPIP_s models, which will be described next.

**Fig. 2.**
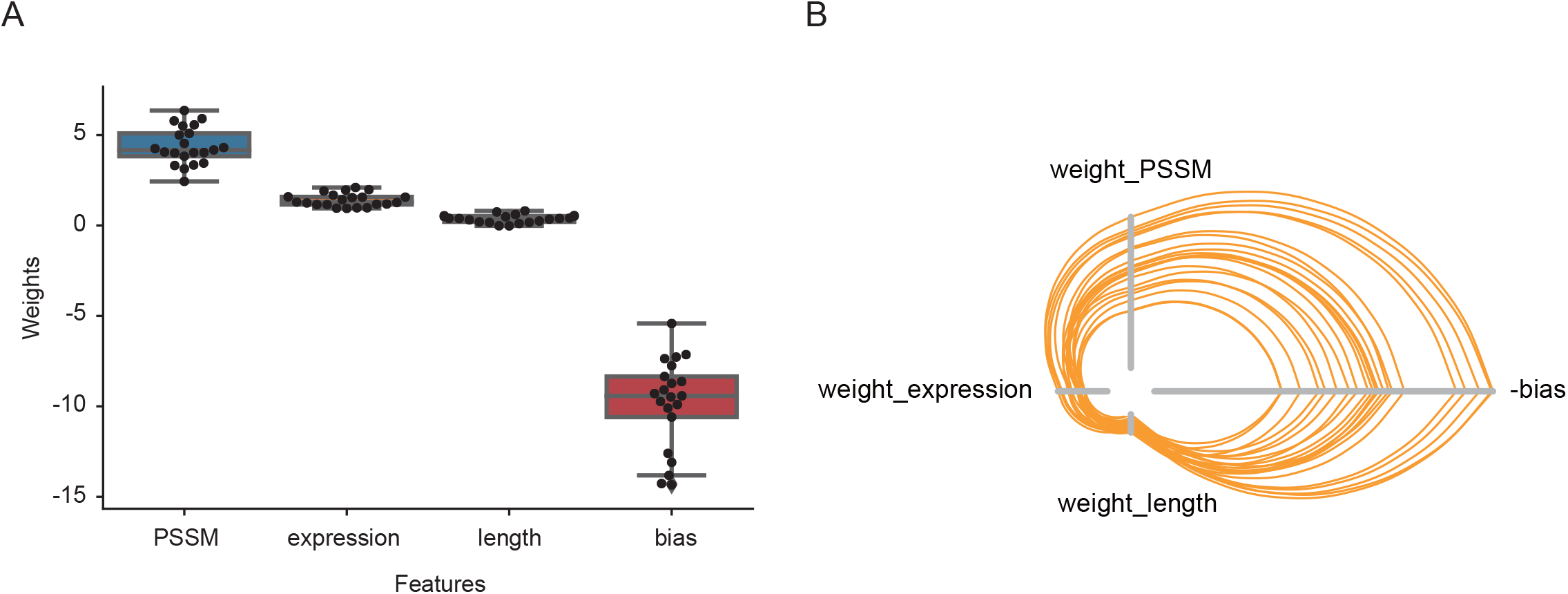
Relationship among weights and biases of 21 EPIP_s models. a) Boxplot for feature weights and biases of EPIP_s models: PSSMs contributed most to the prediction, followed by expressions and peptide lengths. b) Hive plots demonstrating the similar inter-relationships among the feature weights and biases of different EPIP_s models, where the negative biases have been inverted for plotting.

### EPIP and EPIP_s achieved similar performance on epitope presentation prediction

Based on five-fold cross validation, we performed comprehensive evaluations between EPIP and EPIP_s in terms of 0.1% PPV. As shown in Figure 3A, when evaluating across different alleles originating from the same cell line, the performances of EPIP and EPIP_s are quite similar. The mean 0.1% PPV of EPIP is slightly better (0.5741 vs 0.5692), likely due to the more training data available to EPIP, but such a difference is not statistically significant (*P* = 0.1385, paired t-test), and both substantially outperformed predictions based on PSSM scores alone (the mean 0.1% PPV of PSSM is 0.4421).We also performed similar analyses for the same HLA alleles across different cell lines with diverse tissue origins and thus diverse gene expression profiles (Figure 3B). When comparing 0.1% PPV of EPIP and EPIP_s, EPIP is not worse than EPIP_s overall, and even notably better for some cell lines (Figure 3C-F), also likely due to the larger training set of EPIP. Based on these comparisons, we concluded that developing a universal LR model, EPIP, to account for the relationship among these three variables is a valid approach. This seemly simple strategy has substantial practical values, since the EPIP model trained on the selected 21 HLA alleles and cell lines from two different tissue origins can now be straightforwardly extended to previously unseen HLA alleles and tissues without adjusting the model parameters of LR, or requiring extra RNA-seq data for training. We did so for 44 more HLA alleles by incorporating public mono-allele [14] and deconvolved mixed allele [7–15] MS data. We also generated our own MS data to demonstrate as a user case, after evaluating EPIP against other methods.

**Fig. 3.**
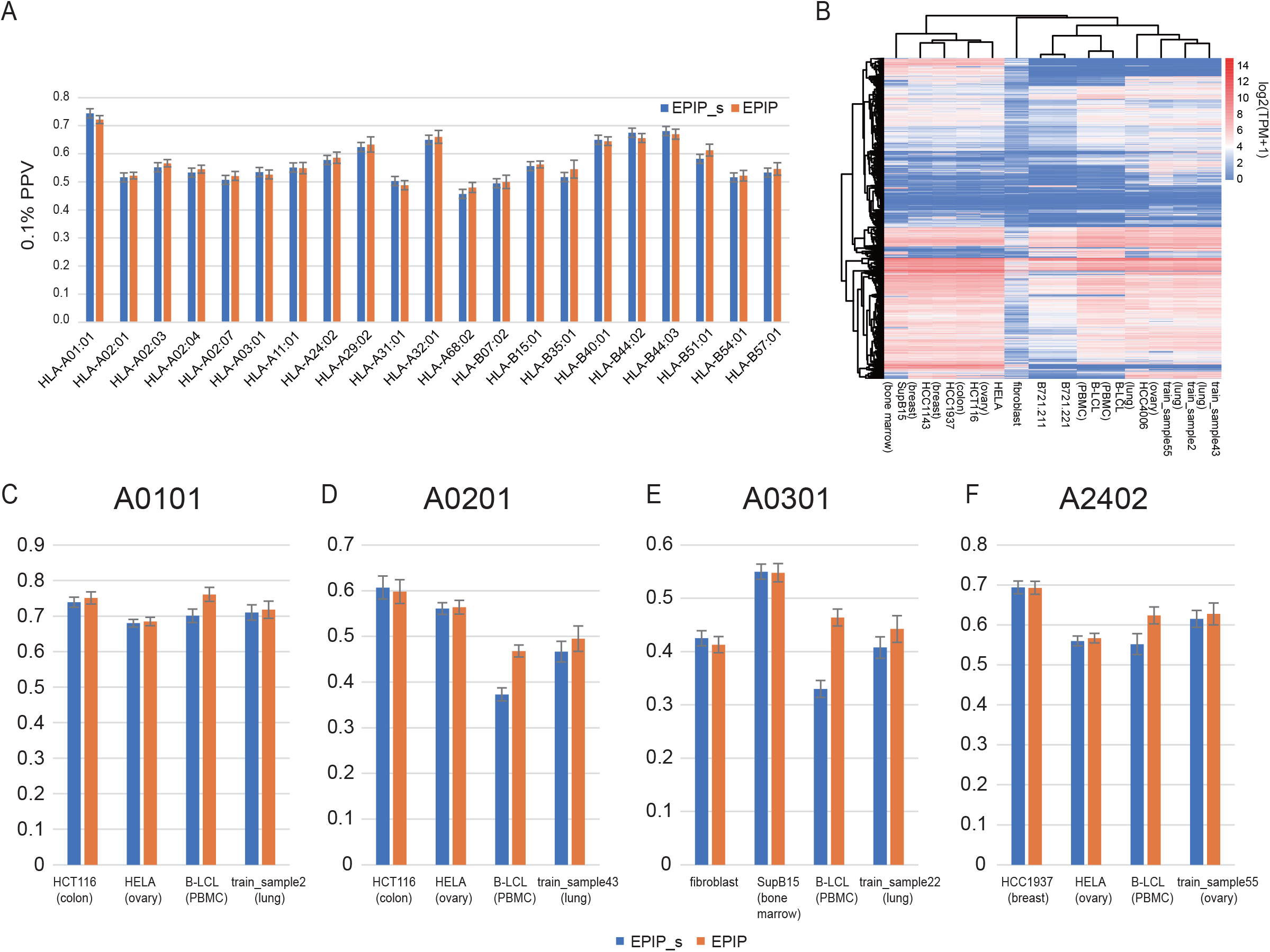
Comparison of prediction performances between EPIP and EPIP_s. a) Performance comparison of EPIP and EPIP_s across different alleles originating from the same B cell line. Error bars indicate the standard deviations across five-fold cross validation. The performance difference between EPIP_s and EPIP across 21 alleles is not statistically significant (*P* = 0.1385, paired t-test) b) Heatmap demonstrates diverse expression profiles of different cell lines originating from different tissues. Each row represents a gene that presents peptides in at least one cell line, a total of 10068 genes was selected. Each column represents a cell line that used in this study, the originated tissue is indicated below the cell line. The expression values are transformed by log2(TPM+1). c-f) Performance comparison of EPIP and EPIP_s for the same allele originating from different cell lines. Error bar indicate the standard deviations across five-fold cross validation.

### EPIP significantly outperforms other existing methods

We evaluated EPIP against MixMHCpred (v2.0), NetMHCpan-4.0 (EL), NetMHCpan-4.0 (BA), and MHCflurry (v1.2.2) on independent validation MS datasets collected from public sources, which contained 29 MS datasets and was not used in the training of EPIP. EPIP significantly outperformed other methods with mean 0.1% PPV of 52.01%, compared to 37.24%, 36.96%, 24.90% and 23.76% achieved by MixMHCpred, NetMHCpan(EL), NetMHCpan (BA) and MHCflurry, with highly significant p-values (*P* = 2.819e-11, 3.960e-11, 1.738e-14, 6.826e-14, respectively). Since other methods in the comparison do not make use of gene expression values, we further combined different gene expression value cutoffs with their predictions. The advantage of EPIP persists and the differences are still highly significant, as shown in Figure 4A. Full comparisons for each validation dataset are provided in Figure S2. The recently developed EDGE method has also been shown to excel at epitope presentation prediction comparing to existing methods, but we were not able to do a fair comparison since it is not publicly available. Nevertheless, we will discuss this further with some qualitative comparisons in the Discussion section to show that the performances of EPIP and EDGE are comparable. Overall, the results on independent MS validation sets showed the clear advantage of EPIP in predicting epitope presentation.

**Fig 4.**
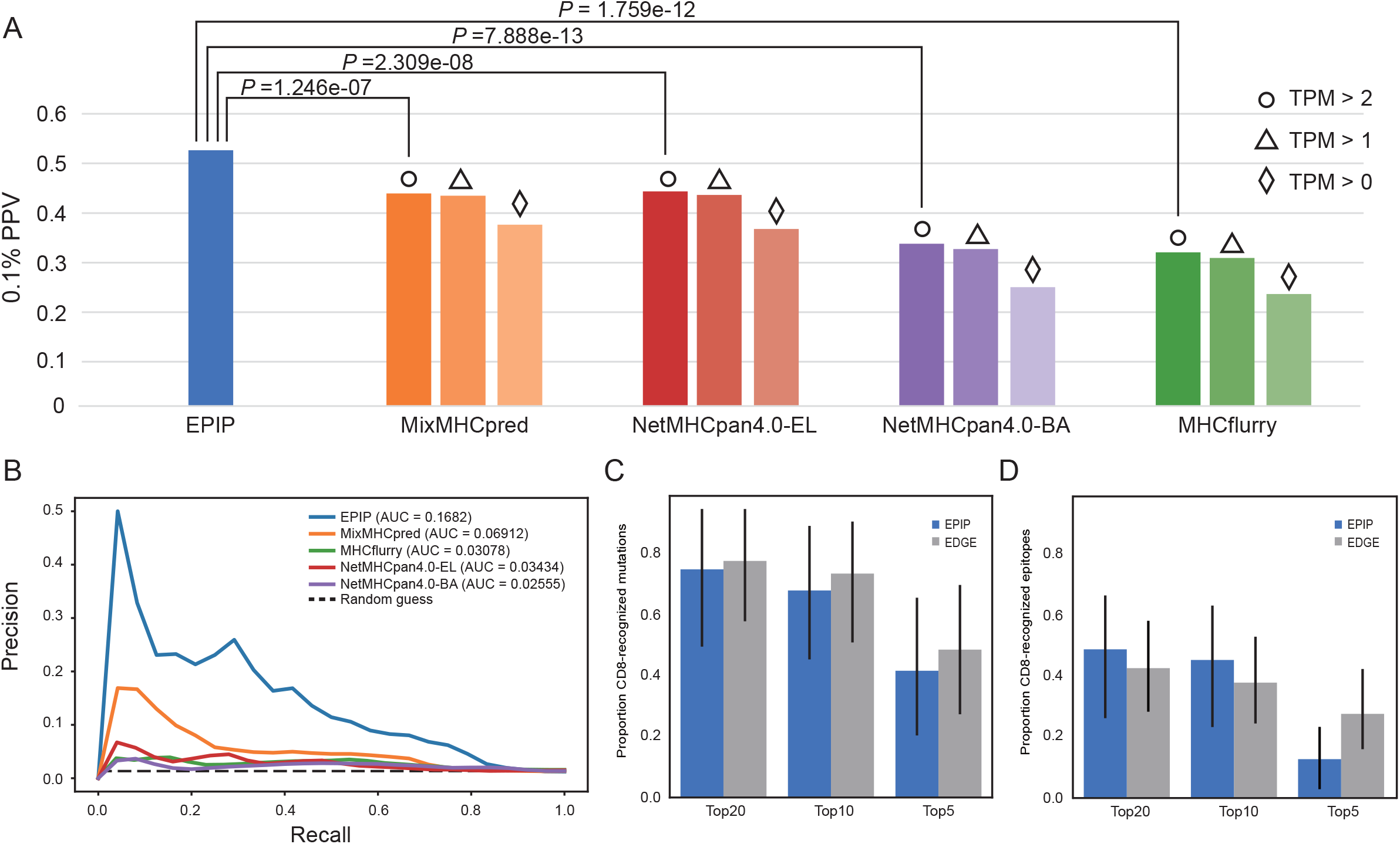
Comparison of prediction performances between EPIP and other methods. a) Performances of EPIP and other publicly available methods on independent MS dataset. The evaluation dataset was obtained from four publications that contained a total of 30 MS datasets, including 16 different tissue origins and 17 HLA alleles. The mean 0.1% PPV of these 30 datasets were displayed. EPIP significantly outperformed other methods with different expression filtering thresholds (TPM > 0, > 1, > 2), and pairwise P-values (paired t-test) for TPM > 2 were shown in the plot. b). Precision-recall curves of EPIP and other publicly available methods on T cell response data. Other methods’ predictions were made with expression threshold of TPM > 0. Comparisons of EPIP and other methods with TPM > 1 and TPM > 2 were displayed in Fig S3. c) Prediction performance of EPIP and EDGE on immunogenic SNVs. EDGE’s performance was recalculated according to the supplementary data 3c of the original paper. d) Prediction performance of EPIP and EDGE on minimal epitopes. EDGE’s performance was taken from Figure 4 of the original paper.

### EPIP is valuable to immunogenicity prediction

For cancer immunotherapies, as well as other clinical applications, the ultimate task is usually to predict the immunogenicity of an epitope. While this remains an elusive goal in general [38], it has recently been shown that accurate prediction of epitope presentation can significantly improve immunogenicity predictions [7]. Therefore, we downloaded the same dataset as used in Bulik-Sullivan et al [7], which was collected from four previous publications [27–30] that consisted of 26 SNVs and 31 neoantigens with pre-existing T-cell responses among 2,023 assayed single nucleotide variants (SNVs) from 17 patients, to evaluate our method against others. We first asked if the increased accuracy of EPIP over other publicly available methods on epitope presentation could translate into better immunogenicity prediction. To quantify this, we plotted the precision recall curves (PRCs) by ranking all SNVs scored by different methods, where the score of a SNV is the maximum score of all possible epitopes generated by the SNV. EPIP’s area under PRC (AUPRC) was more than twice that of other methods, and kept the advantages when we applied expression value threshold of 1 or 2 TPM to other methods’ predictions (Figure 4B and Figure S3). We also compared EPIP with EDGE at the level of ranking mutations and minimal epitopes (the recognized 8-11-mers overlapping the mutation) as originally done in Bulik-Sullivan et al [7]. EPIP is very comparably to EDGE on these two evaluations, especially for the top 10 and top 20 ranked mutations and epitopes. When prioritizing the mutations, the number of immunogenic SNVs ranked in the top 20, 10, and 5 were 18 (69.23%), 16 (61.54%) and 11 (42.31%) for EPIP, and 19 (73.08%), 18 (69.23%) and 12 (46.15%) for EDGE (Figure 4C). When prioritizing the minimal epitopes, EPIP ranked 15 (48.39%), 12 (38.71%) and 5 (16.13%) CD8+ recognized epitopes in the top 20, 10 and 5. Therefore, despite being a much simpler model trained with less data comparing to EDGE, EPIP is also quite valuable for immunogenicity prediction in neoantigen-based cancer immunotherapies and is a significant improvement over other publicly available methods.

### EPIP can be extended to support more alleles by incorporating new MS data

Due to its simplicity and modular structure, EPIP can be straightforwardly extended to support more HLA alleles by incorporating new MS data. An allele that we are particularly interested in is HLA-A*33:03, which appears often in the Chinese population but with very limited MS or binding affinity data prior to this publication (more details in the supplementary material). We first performed MS profiling of eluted HLA ligands on the HCC4006 lung cancer cell line, which is homozygous on all three HLA alleles (HLA-A*33:03, HLA-B*44:03 and HLA-C*07:06, respectively). Upon deconvolution of MS data by GibbsCluster and removing the trash cluster, we were only able to obtain two clusters, likely due to the low expression of HLA-C alleles. We were able to assign one of the clusters to HLA-B*44:03 by comparing to the known motif of HLA-B*44:03 obtained from mono-allelic HLA-B*44:03 cell line [12]. We believe the other cluster should belong to HLA-A*33:03, since the motif is quite similar to the A33:03 motif predicted by NetMHCpan-4.0 in its pan-specific mode (although binding affinity has not been directly profiled for this allele) and ~80% of the peptides in this cluster has binding affinity < 500nm to HLA-A*33:03 as predicted by NetMHCpan-4.0. To further check on this, we also performed t-SNE [39] analysis for the peptides in this cluster, together with peptides of HLA-*A02:01 and HLA-A*11:01 in EPIP’s training set. As shown in Figure S4, peptides from this cluster is much closer to those from HLA-A*11:01 than HLA-A*02:01, consistent with the fact that HLA-A*33:03 and HLA-A*11:01 belong to the same HLA supertype while HLA-A*02:01 belongs to a different supertype. Based on these evidences, we confidently assigned this cluster to HLA-A*33:03. By inputting these HLA-A*33:03-specific peptides to EPIP, where a dedicated module has been built to accept new allele-specific MS data, it can learn the corresponding PSSM and length distribution and add HLA-A*33:03 to its supported alleles. The overall work flow is illustrated in Figure 5.

**Fig 5.**
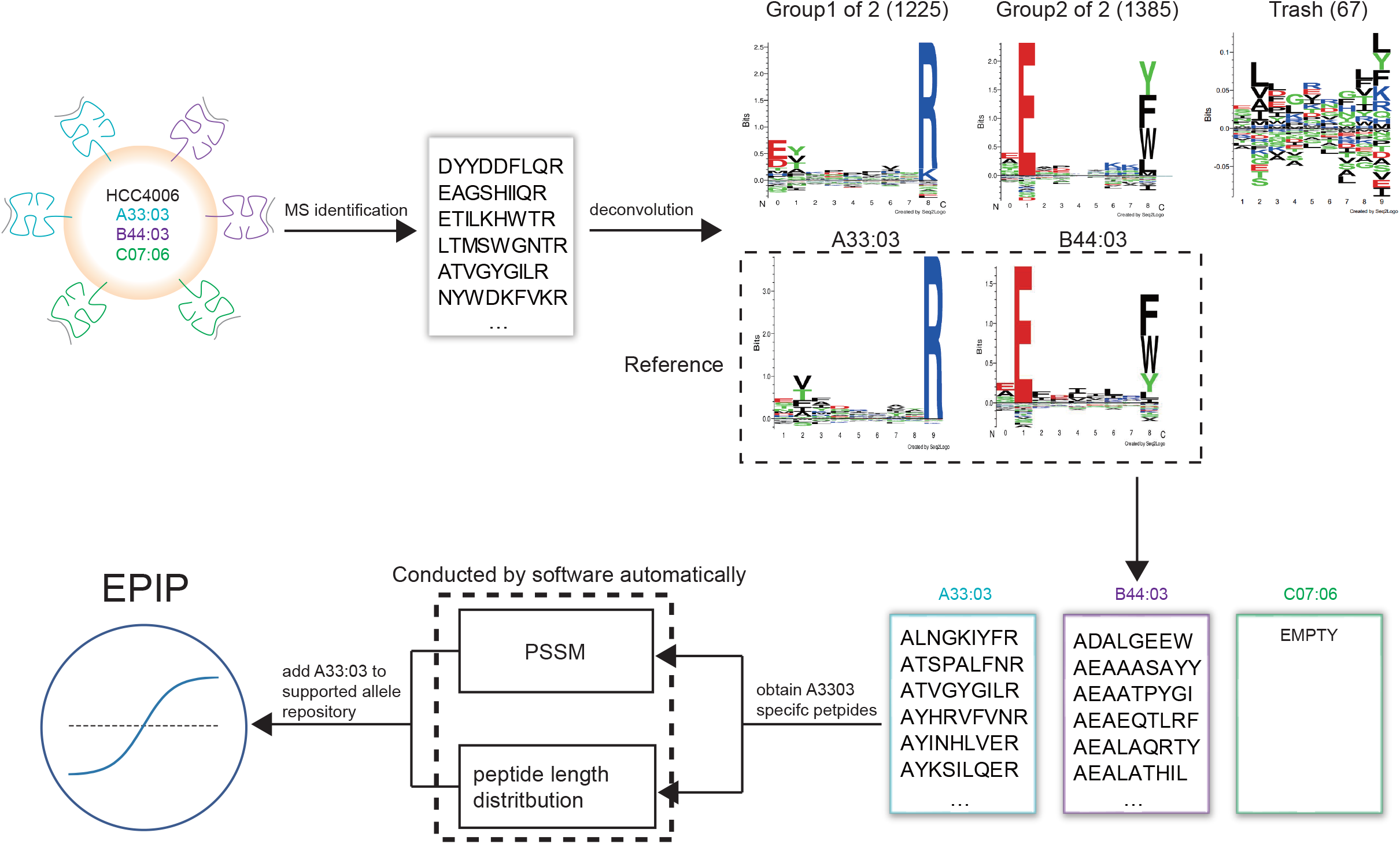
The workflow of adding new allele HLA-A*33:03 to EPIP’s supported allele repository. HCC4006 is homozygous on all three HLA alleles (HLA-A*33:03, HLA-B*44:03 and HLA-C*07:06, respectively). Presented peptides from HCC4006 were identified by MS (see Methods). The derived peptidome was then deconvolved using GibbsCluster2.0. Apart from a trash cluster of 67 peptides, two peptide clusters were identified. One cluster matched the sequence motif of B4403 (derived from HLA-B*44:03 mono-allelic dataset [12]), and the other matched the sequence motif of A33:03 derived from NetMHCpan4.0 online motif reviewer. Noted that due to the lack of training data, NetMHCpan4.0 generated the A33:03 motif based on training data from other HLA alleles in a pan-specific mode, which could explain the slight difference between the motif we discovered and the NetMHCpan-4.0 provided. We then inputted the HLA-A*33:03-specific peptides to EPIP, which could learn PSSM and length distributions automatically, and resulted in an updated version of EPIP that could support the newly added allele.

## Discussions

Besides making use of recently available, large-scale MS immunopeptidome data, our method clearly demonstrated the advantage of integrating gene expression levels in a principled way when modelling epitope presentation, but the use of gene expression levels could present its own challenges. Even when discovering neoantigens for cancer patients where RNA-seq is routinely preformed, this information could be missing or inaccurate. In this case, since it is well-known that gene expression levels are largely preserved in the same tissue [40], one could make use of the gene expression levels quantified in a different sample from a similar tissue origin. For example, we showed that when replacing the expression levels of train_sample55 with HELA, which are both originated from ovary tumors, the performance of EPIP on HLA-A*2402 only dropped a little and still significantly outperformed other public methods (Figure S6). In other applications of epitope prediction, such as those originating from pathogens, the definition of gene expression levels could be problematic, but since virus employ the human cell machinery to produce their protein, the virus epitopes may share the same processing disciplines with human epitopes. As more sophisticated RNA-seq analysis methods are being developed to analyze the transcriptome of virus infected cells and calculate the TPM/RPKM of specific virus proteins [41], EPIP could be used to predict virus epitopes in these cells as well.

The incorporation of gene expression levels in EPIP is similar in principle to what has been done by EDGE [7]. Although EDGE is neither publicly available nor straightforward to be reproduced, some comparisons with EDGE on epitope presentation prediction would still be informative. To achieve this, we used the same evaluation criteria as in EDGE to compare against other affinity-based methods, i.e., to further reduce the positive to negative ratio, or prevalence, to 1:2,500 and 1:10,000 and checked the PPVs of EPIP at 40% recall. Although it is difficult to justify the optimal threshold of prevalence for neoantigen prediction, further reducing this threshold does have the side effect of enlarging the performance gap between EPIP and other methods. As shown in Figure S5, EPIP reached a more than 10-fold improvement over MHCflurry with 1:10,000 prevalence, similar to what has been achieved by EDGE. Combined with the fact that EPIP and EDGE performed similarly on the same immunogenicity test data, we concluded that the performances of EPIP and EDGE should be quite comparable. Since EPIP is a much simpler model comparing to EDGE, this also implies that the performance gain of EDGE is to a larger degree contributed by its large-scale training data than the more sophisticated deep learning framework.

One clear future work for EPIP is to add more allele support as more MS data becomes available, both from public sources and our own experiments. Enabled by EPIP’s simple and modular structure, doing so is quite straightforward as demonstrated by adding the support of HLA-A*33:03, and we will keep doing so and update our web tool over time. We chose to adopt simple machine learning techniques, i.e., PSSMs and LR, for this initial version of EPIP for easy analysis and development. More recent deep learning techniques for sequence analysis, mostly developed in the field of natural language processing [42, 43], can make better use of large datasets and model different contributing factors more effectively, and could lead to more powerful variable length, pan-specific models. This is another promising direction for future versions of EPIP. Moreover, immunogenicity prediction in a clinical setting depends on more than epitope presentation [44–46], and we believe it is important to account for the extra factors contributing to immunogenicity as well. To that end, community efforts to generate more immunogenicity data and improved experimental techniques to profile TCR and peptide-MHC interactions [47, 48] might lead to larger datasets and deeper biological understanding that can eventually enable accurate predictions of immunogenicity.

## Conclusion

In this paper, we have developed a user-friendly, publicly available epitope prediction tool, EPIP, that incorporates information from both MS and RNA-seq data. It significantly outperforms other publicly available tools and can be easily extended to support more HLA alleles when new MS data becomes available. We believe EPIP is a timely contribution to the community to advance cancer immunotherapies and beyond.

## List of abbreviations

HLA: human leukocyte antigen
MS: Mass spectrometry
PSSMs: position score specific matrices
LR: logistic regression
EPIP: Epitope Presentation Integrated Prediction
MHC: major histocompatibility complex
TCRs: T-cell receptors
PMF: probability mass function
PPV: positive predictive value
TMG: tandem minigene
PRCs: precision recall curves

## Declarations

### Ethics approval and consent to participate

Not applicable

### Consent for publication

Not applicable

### Availability of data and materials

The MS data of HCC4006 generated during the current study are available in the CNGB Nucleotide Sequence Archive (CNSA: https://db.cngb.org).

### Authors’ contributions

SQ, LJL, WH, BL designed the project. WH, XH, SZ, GL, JL implemented the EPIP model. WH and YL collected the data for training and evaluation of the model. XL, LC, SL cultured the HCC4006 cell line. XL, LC, SL, GH, ZL, YR generated the MS data of HCC4006. LZ, YL, WH, WL, HX, ZR analyzed the MS data of HCC4006. LJL, WH, SQ, YL, XL, LZ, BL wrote the manuscript. LJL, SQ, BL supervised the project. All authors read and approved the final manuscript.

## Acknowledgements

We wish to thank Dian Dong for helping with formatting the paper and references.

## Footnotes

1 The number of immunogenic epitopes is the same as in a more recent publication [31] retrieving data from the same source.

