## Supplementary materials for "EPIP: MHC-I epitope prediction integrating mass spectrometry derived motifs and tissue-specific expression profiles"

**Stacking method for building PSSM**

To build PSSM, A possible way is to use part of the MS peptides to build PSSM and score the remaining data to train EPIP_s. However, this method may lead to not enough data to build PSSM or train EPIP_s, thus mitigate the performance. Another way is to use all the MS peptides to build PSSM and use PSSM to score all the training peptides, then use the scored training peptides to train EPIP_s. The potential problem of the method is that the MS peptides are trained in both PSSM and EPIP_s, which may cause overfitting. To fully utilize the data and avoid overfitting, we used the stacking method [25] to build PSSM. Specifically, the MS peptides were split into five folds for iteration. During each iteration, a fold of data was held out and the other four folds were used to build PSSM. The generated PSSM was used to calculate the PSSM score for the held-out data. The scored training dataset was obtained after the iteration. Since the scored peptides were not used to train PSSM, the overfitting problem was avoided. To calculate PSSM score for the decoy peptides, PSSM generated from each iteration was used to score the decoy peptides and the averaged PSSM score across all the PSSMs were used as the final score. The stacking scheme is illustrated in Fig S1.

**Sequence logo plots**

The reference sequence logo for the clusters identified by Gibbscluster (Fig 5) is generated using Seq2Logo-2.1 [49].

**EPIP independent evaluation**

Peptides from fibroblast, HCT116, HCC1143, HCC1937, SupB15 were collected from Bassani-Sternberg et al [11]. Peptides from B-LCL were collected from Pearson et al [13]. Peptides from train_sample2, train_sample3, train_sample17, train_sample22, train_sample26, train_sample40, train_sample43, train_sample55, train_sample58 were collected from Bulik-Sullivan [7]. Because these samples expressed multiple alleles, we assigned the peptides to our interested allele using Gibbscluster [24]. Peptides from HeLa cell line were collected from Trolle et al [16], which was a soluble HLA single-allele mass spectrometry dataset thus no need to perform deconvolution. 999-fold decoy peptides were added to the MS peptide dataset for each cell line to construct the final test dataset. Because the Bassani-Sternberg data and the Trolle data did not have associated expression data, the expression data of cell lines from these studies were downloaded from ENCODE [50] and TRON Cell Line Portal [51], where expression values from the latter were represented as RPKM and transformed to TPM [51]. RNA-seq data provided by Pearson et al [13] is used to recalculate the expression profile of B-LCL as described in Method. The expression profile provided by Bulik-Sullivan [7] was used directly since RNA-seq data was not available. This independent dataset was used to evaluate EPIP, NetMHCpan4.0-EL, NetMHCpan4.0-BA, MixMHCpred and MHCflurry, and the 0.1%PPV was compared to each other. To demonstrate EPIP is comparable to EDGE, we evaluate PPV at 40% recall for each software using A0201 peptides from HeLa, A2402 peptides from HCC1937 and B4402 peptides from B-LCL (Fig S5).

**MS data generation and processing for HCC4006 cell line**

**HLA-I peptidomes sample preparation**

HLA-I peptidomes were obtained from HCC4006 cell line as described previously [11]. In brief, 1E9 cell pellets were dissociated using 40 ml of lysis buffer with 0.25% Sodium deoxycholate, 1% n-octyl glucoside, 100 mM PMSF and protease inhibitors cocktail in PBS at 4 °C for 60 min. Lysate were further cleared by 30 min centrifugation at 14,000 g. Cleared lysate were immunoaffinity purified with pan-HLA class I complexes antibody covalently bound to Protein-A Sepharose CL-4B beads. Beads were then washed first with 10 column volumes of 150 mM NaCl, 20 mM Tris HCl (buffer A), 10 column volumes of 400 mM NaCl, 20 mM Tris HCl, 10 volumes of buffer A again, and finally with 10 column volumes of 20 mM Tris HCl, pH 8.0. The HLA-I molecules were eluted at room temperature using 0.1 N acetic acid. Eluate were then loaded on Sep-Pak tC18 cartridges (Waters, 50mg) and wash with 0.1% TFA. The peptides were separated from HLA-I complexes on the C18 cartridges by eluting with 30% ACN in 0.1% TFA and concentrated to 20 $\mu$l using vacuum centrifugation. Finally, 5 $\mu$l sample was used for MS analysis.

**HLA-I Peptides sequencing by LC-MS/MS**

HLA peptides were separated by a nanoflow HPLC (15 cm long, 75 $\mu$m inner diameter column with ReproSil-Pur C18-AQ 1.9 $\mu$m resin) and coupled on-line to a Fusion Lumos mass spectrometer (Proxeon Biosystems, Thermo Fisher Scientific) with a nanoelectrospray ion source (Proxeon Biosystems). Peptides were eluted with a linear gradient of 2–30% buffer B (80% ACN and 0.5% acetic acid) at a flow rate of 250 nl/min over 65 min. Data was acquired using a data-dependent "top 10" method, which isolated and fragmented them by higher energy collisional dissociation (HCD). Full scan MS spectra were acquired at a resolution of 120,000 at 350-1500 m/z with a target value of 3e6 ions. The ten most intense ions were isolated and accumulated to an AGC target value of 1e5 with a maximum injection time of 50 ms. The peptide match option was disabled. MS/MS resolution was 60,000 at 100 m/z. The interpretation of MS data is described in Methods.

**Peptide distance calculation and visualization, related to Fig S4**

The major procedures are conducted as described by Abelin et al [12]. The A02:01 and the A11:01 specific peptides are from EPIP's training set. The entropy at each 9mer position was first calculated using MolecularEntropy() function from HDMD R package for A02:01, A11:01 and the potential A33:03 peptidome. Then all the peptides were pooled together and entropies across three peptidome were averaged. Next, we employed the following function to compute the distance between every pair of 9mer peptides in the peptide pool:

$$d(s_1, s_2) = \frac{1}{9}\sum_{i=1}^{9} distPMBEC(s_{1i}, s_{2i}) * (1 - entropy_i)$$

In the function, $s_1$ and $s_2$ mean two different peptide sequences, $PMBEC$ is an amino acid similarity matrix derived from the binding affinity data [54], and $distPMBEC$ , defined as $max(PMBEC) - PMBEC$, is a 20x20 matrix capturing residue dissimilarities. $s_{1i}$ and $s_{2i}$ mean the amino acid at position $i$ of the peptide sequences. $entropy_i$ means entropy at position $i$ of the average entropy. The derived pairwise peptide distance matrix was then visualized using t-SNE from scikit-learn 0.19.0 [26].

**Motif distance calculation**

The motif distances are calculated as described by Bassani-Sternberg et al [52]. Briefly, 9mer peptides in each motif were selected to build PWMs with pseudo count 1. Then, Euclidean distance was calculated between two PWMs:

$$D^2 = \frac{1}{9}\sum_{i=1}^{9}\sum_{a=1}^{20}(M_{ia} - M'_{ia})^2$$

In the function, $M_{ia}$ and $M'_{ia}$ mean PWM value of amino acid a at positon i.

**Cysteine bias examination**

Although MS data is considered to show a more accurate view of peptide presentation comparing with binding affinity data, MS analyzed peptides also have its own biases. One of the known limitations is the low frequency of cysteine observed in MS peptidome data, as cysteine is prone to post-translational modifications that are typically not included in database searches [52]. However, the benefits brought by MS peptidome training set are greater than the disadvantages, even in predictions of cysteine-containing T cell epitopes, as demonstrated in EDGE [7]: while binding affinity methods only ranked zero or one out of seven cysteine-containing T cell-recognized epitopes in the top five, EDGE can rank three out of seven. We performed similar analysis and our EPIP method can even rank six out of seven cysteine-containing epitopes in the top five. We also explored whether cysteine frequency adjusted PSSMs could further improve prediction on cysteine-containing epitopes, and the adjustments are detailed below.

We defined PSSM as a matrix of M rows (Amino acid; M=20) and N columns (Length; N=9). Each element $P_{ai}$ in the matrix is the likelihood of a given character (amino acid) at its position. We calculated the element $P_{ai}$ through

$$P_{a_i} = log\frac{F_{a_i}+\omega}{BG_a} \quad (1),$$

where $F_{ai}$ denotes the frequency of amino acid $a$ at position i, $BG_a$ denotes the background frequency of amino acid $a$ from uniprot database [53], and ω is a random value (ranging from 0 to 1) generated from Dirichlet distribution [54]. Normally, the $BG_a$ of cysteine is 0.013685033. In modified PSSM_1, we replaced the $BG_a$ of cysteine with the cysteine frequency at nonanchor positions in peptidomics training data. In modified PSSM_2, we used equation (2) for cysteine but equation (1) for other amino acids.

$$P_{a_i} = log\frac{F_{a_i}+\omega+0.012232033}{BG_a} \quad (2),$$

where 0.012232033 is the difference between cysteine frequency at nonanchor positions in peptidomics and in human proteome.

As shown in Table S2, by using the two modified PSSMs, EPIP was still not able to predict all seven cysteine-containing epitopes, but it performances on cysteine-free HLA ligands predictions declined considerably. Therefore, cysteine adjusted PSSMs were not used in the final version of EPIP.

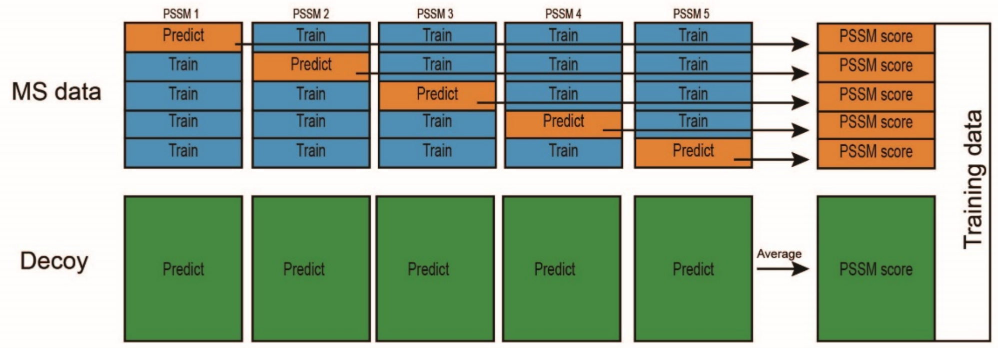


Fig S1. Stacking scheme of building PSSM.

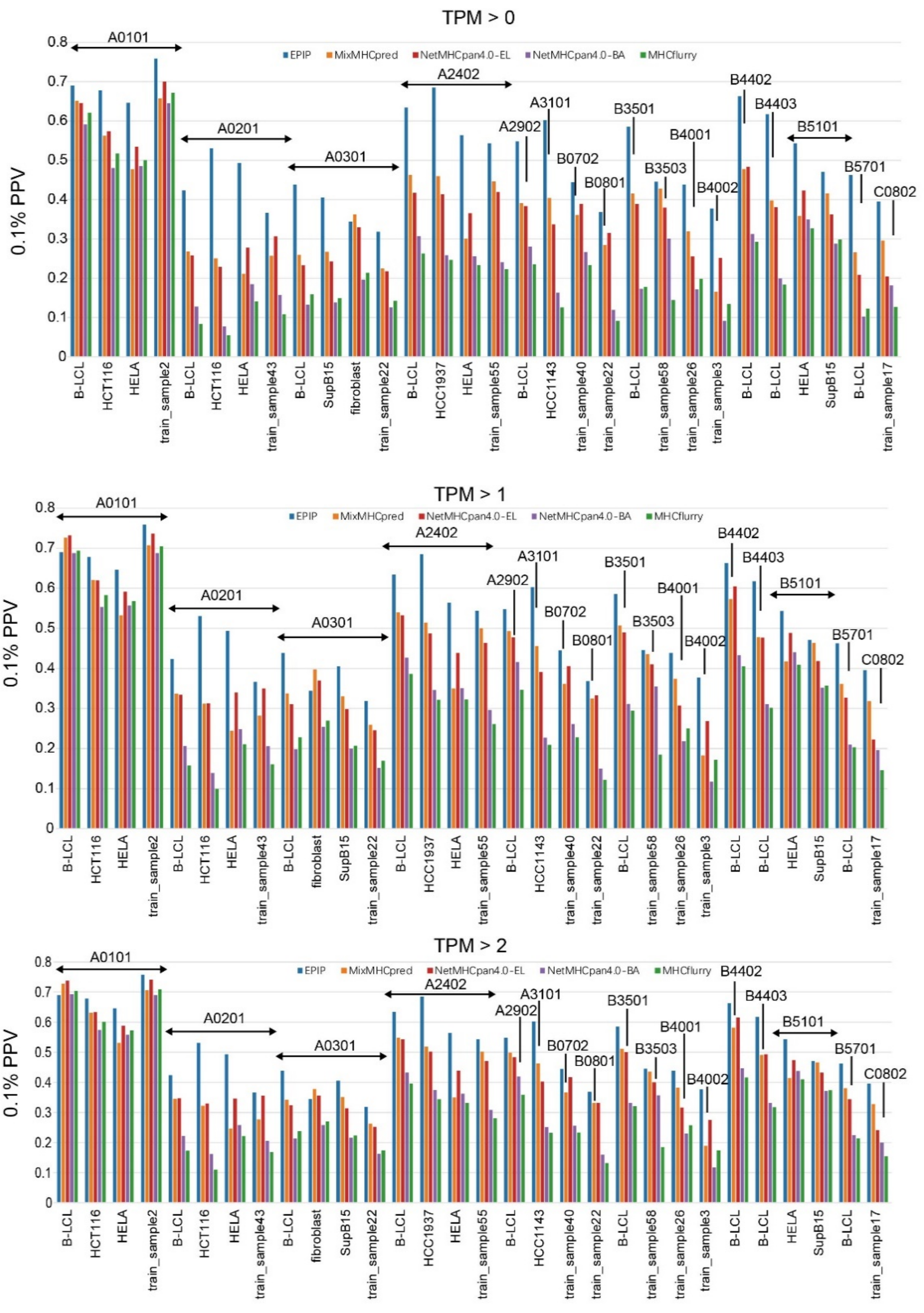


Fig S2, related to Fig. 4 A. Evaluation of EPIP on MS dataset against other software. MS dataset is collected from 4 published datasets [7,11,13,16]. HELA is a mono-allelic cell line, so the MS data is used directly. For other cell lines, the MS data is deconvolved first to obtain peptides that belong to interested allele. Then each MS dataset is combined with 999-fold decoy peptides that excluded the MS peptides. The performance of EPIP was compared with that of MixMHCpred, NetMHCpan4.0-EL, NetMHCpan4.0-BA, MHCflurry combined with TPM > 2, 1, 0.

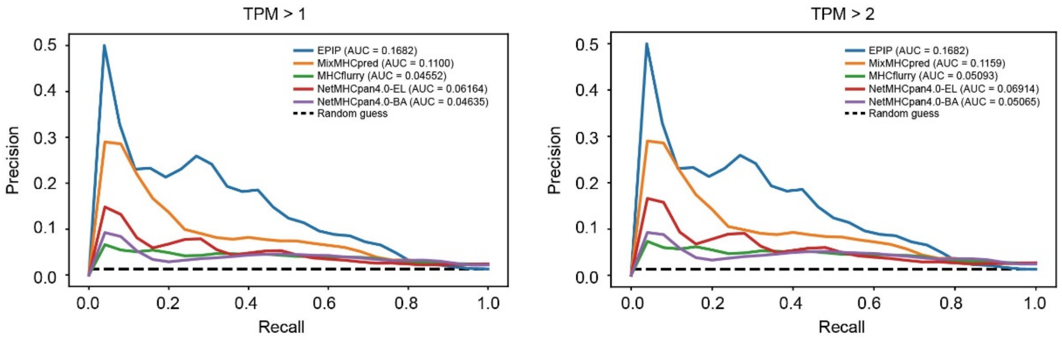


Fig S3, related to Fig. 4 B. Evaluation of EPIP on T-cells recognized SNVs dataset against other software. Although the performance of other software has obvious improved after applying additional expression value cutoffs (compared with TPM > 0, as shown in Fig 4. B), EPIP still shows significant advantage. Besides, the improvement of performance is limited when TPM cutoff changed from 1 to 2 compared with that changed from 0 to 1.

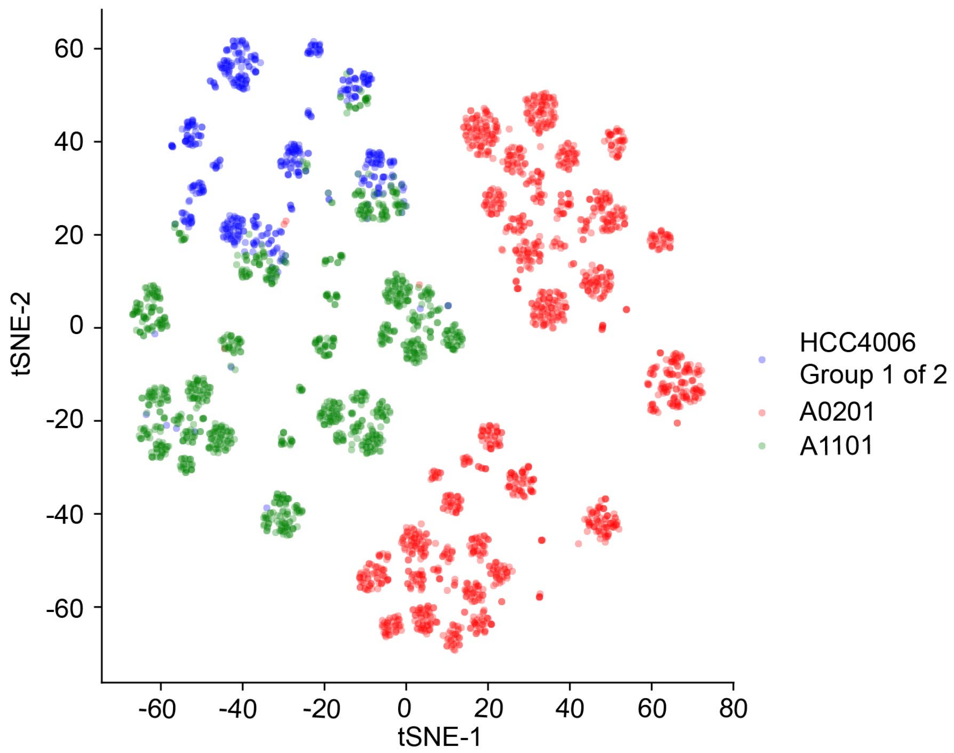


Fig S4. t-SNE displaying the peptidome distance between the first group deconvolved peptides from HCC4006 and MS peptides of A11:01 as well as A02:01. Since A11:01 and A33:03 belong to the same supertype, to verify if we can assign the uncertain motif from deconvolution to A33:03, we calculate the distance matrix of MS peptides of A1101 and the unknown motif and visualize using t-SNE. We also conduct the same analysis using MS peptides of A0201 as control group. It shows a clear division between the distribution of MS peptides for A1101 and A0201, while the distribution of the peptides from the uncertain motif overlaps with that of A1101. Thus, we are able to confidently assign the uncertain motif to A33:03.

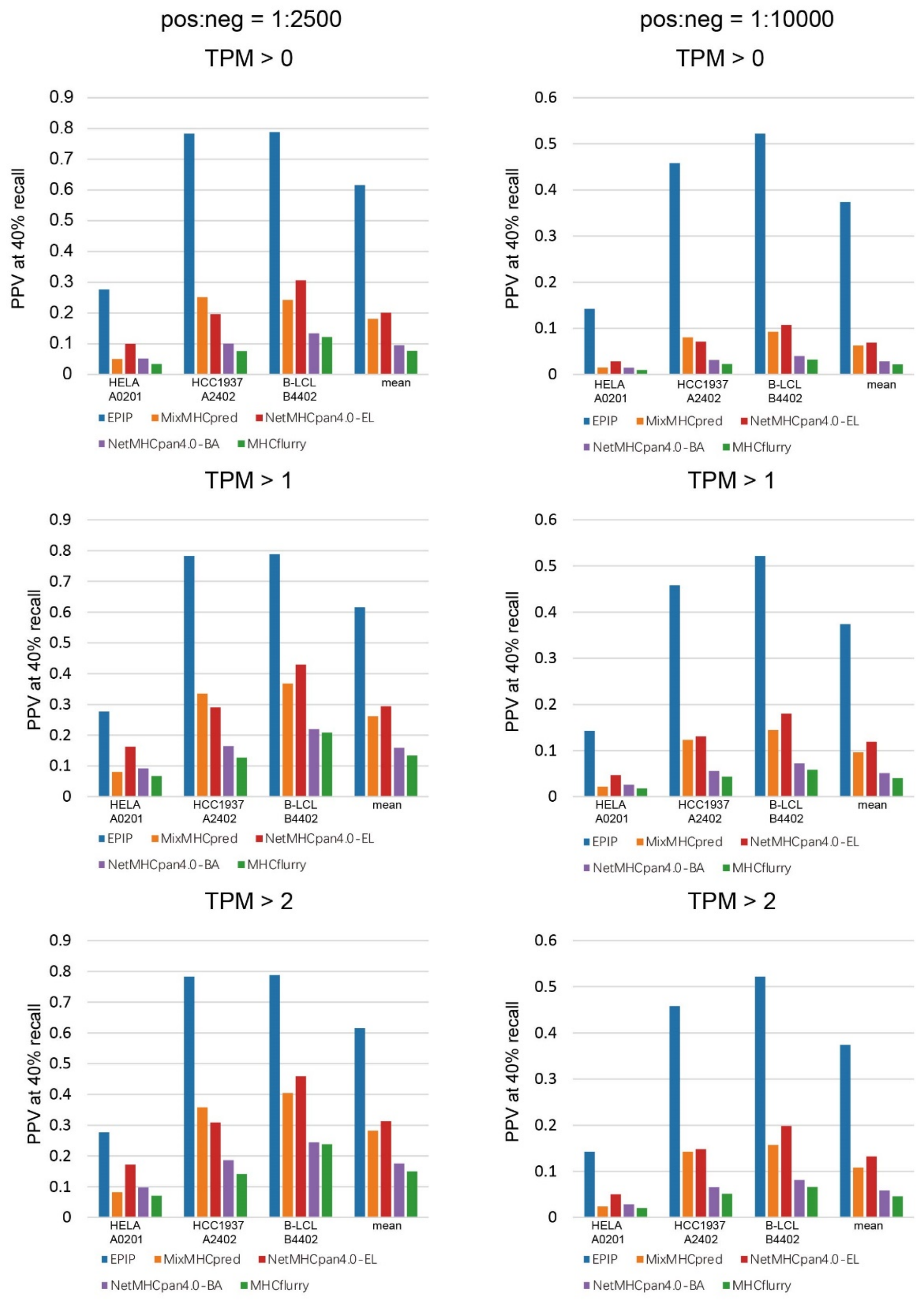


Fig S5. Evaluating EPIP against other methods. As EDGE is not available, to prove EPIP is comparable to EDGE, we evaluated the PPV at 40% recall of EPIP and other software on the test dataset in which the ratio of positive and negative dataset is 1:2500 and 1:10000, which was the approach used by EDGE for independent evaluation against MHCflurry. The advantage of EPIP achieved a close to 10-fold improvement over MHCflurry with 1:2,500 prevalence and the improvement is more than 10-fold with 1:10,000 prevalence.

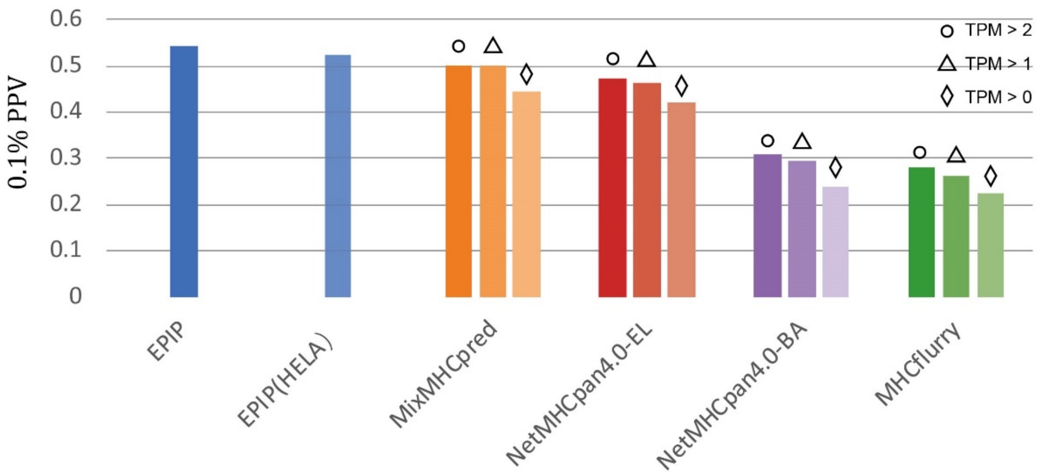


Fig S6. Evaluation of EPIP using MS data of train_sample55. When replacing the expression levels of train_sample55 with HELA's, the performance of EPIP (labeled as EPIP(HELA)) on predicting HLA-A*2402 epitopes only dropped a little compared with using train_sample55's expression levels (labeled as EPIP), and still outperformed that of MixMHCpred, NetMHCpan4.0-EL, NetMHCpan4.0-BA, MHCflurry combined with hard-filtering threshold TPM > 2, 1, 0.

Table S1 Training data of EPIP.

| Allele | Length: Peptide number | | | Source | Type |
|---|---|---|---|---|---|
| A0101 | 9:437 | 10:325 | 11:218 | Abelin2017 | mono-allele |
| A0201 | 9:1652 | 10:391 | 11:407 | Abelin2017 | mono-allele |
| A0203 | 9:1236 | 10:454 | 11:90 | Abelin2017 | mono-allele |
| A0204 | 9:1407 | 10:166 | 11:153 | Abelin2017 | mono-allele |
| A0207 | 9:2394 | 10:430 | 11:382 | Abelin2017 | mono-allele |
| A0301 | 9:765 | 10:388 | 11:307 | Abelin2017 | mono-allele |
| A2402 | 9:1412 | 10:410 | 11:310 | Abelin2017 | mono-allele |
| A2902 | 9:566 | 10:159 | 11:87 | Abelin2017 | mono-allele |
| A3101 | 9:425 | 10:194 | 11:294 | Abelin2017 | mono-allele |
| A6802 | 9:913 | 10:360 | 11:243 | Abelin2017 | mono-allele |
| B3501 | 9:514 | 10:142 | 11:111 | Abelin2017 | mono-allele |
| B4402 | 9:412 | 10:329 | 11:206 | Abelin2017 | mono-allele |
| B4403 | 9:368 | 10:274 | 11:153 | Abelin2017 | mono-allele |
| B5101 | 8:469 | 9:768 | 10:77 | Abelin2017 | mono-allele |
| B5401 | 9:627 | 10:196 | 11:146 | Abelin2017 | mono-allele |
| B5701 | 9:481 | 10:305 | 11:219 | Abelin2017 | mono-allele |
| A1101 | 9:1191 | 10:558 | 11:365 | Pearson2016 | deconvolution |
| A3201 | 9:439 | 10:78 | 11:67 | Pearson2016 | deconvolution |
| B0702 | 9:2048 | 10:703 | 11:411 | Pearson2016 | deconvolution |
| B1501 | 9:1096 | 10:422 | 11:166 | Pearson2016 | deconvolution |
| B4001 | 9:801 | 10:366 | 11:154 | Pearson2016 | deconvolution |
| C0102 | 8:102 | 9:987 | 10:235 | Di Marco2017 | mono-allele |
| C0202 | 8:116 | 9:1533 | 10:214 | Di Marco2017 | mono-allele |
| C0303 | 8:91 | 9:852 | 10:99 | Di Marco2017 | mono-allele |
| C0304 | 8:251 | 9:1601 | 10:176 | Di Marco2017 | mono-allele |
| C0401 | 8:467 | 9:1161 | 10:153 | Di Marco2017 | mono-allele |
| C0501 | 8:626 | 9:1563 | 10:249 | Di Marco2017 | mono-allele |
| C0602 | 8:47 | 9:870 | 10:32 | Di Marco2017 | mono-allele |
| C0701 | 8:55 | 9:310 | 10:19 | Di Marco2017 | mono-allele |
| C0702 | 8:116 | 9:589 | 10:53 | Di Marco2017 | mono-allele |
| C0802 | 8:792 | 9:2231 | 10:330 | Di Marco2017 | mono-allele |
| C1203 | 8:146 | 9:1160 | 10:53 | Di Marco2017 | mono-allele |
| C1402 | 8:484 | 9:1604 | 10:313 | Di Marco2017 | mono-allele |
| C1502 | 8:191 | 9:1639 | 10:56 | Di Marco2017 | mono-allele |
| C1601 | 8:685 | 9:1899 | 10:106 | Di Marco2017 | mono-allele |
| C1701 | 8:120 | 9:418 | 10:49 | Di Marco2017 | mono-allele |
| A2301 | 9:627 | 10:145 | 11:92 | SysteMHC | database |
| A2501 | 9:419 | 10:296 | 11:59 | SysteMHC | database |
| A2601 | 9:240 | 10:88 | 11:35 | SysteMHC | database |
| A2901 | 8:6 | 9:153 | 10:9 | SysteMHC | database |
| A3001 | 9:35 | 10:19 | 11:12 | SysteMHC | database |

| | | | | | |
|---|---|---|---|---|---|
| A3004 | 9:77 | 10:15 | 11:22 | SysteMHC | database |
| A6801 | 9:3305 | 10:2031 | 11:1486 | SysteMHC | database |
| B0801 | 8:1418 | 9:2292 | 10:79 | SysteMHC | database |
| B1301 | 8:8 | 9:21 | 10:0 | SysteMHC | database |
| B1402 | 8:36 | 9:161 | 10:7 | SysteMHC | database |
| B1511 | 8:2 | 9:34 | 10:8 | SysteMHC | database |
| B15186 | 8:223 | 9:975 | 10:165 | SysteMHC | database |
| B1801 | 8:929 | 9:1103 | 10:140 | SysteMHC | database |
| B1803 | 8:44 | 9:67 | 10:1 | SysteMHC | database |
| B2701 | 9:326 | 10:138 | 11:69 | SysteMHC | database |
| B2705 | 9:2903 | 10:1529 | 11:936 | SysteMHC | database |
| B3503 | 9:1184 | 10:335 | 11:278 | SysteMHC | database |
| B3508 | 9:507 | 10:125 | 11:57 | SysteMHC | database |
| B3701 | 8:92 | 9:265 | 10:9 | SysteMHC | database |
| B3801 | 9:106 | 10:20 | 11:7 | SysteMHC | database |
| B3901 | 8:150 | 9:764 | 10:115 | SysteMHC | database |
| B3906 | 8:529 | 9:681 | 10:169 | SysteMHC | database |
| B3924 | 8:133 | 9:156 | 10:12 | SysteMHC | database |
| B4002 | 9:1901 | 10:878 | 11:523 | SysteMHC | database |
| B4101 | 9:396 | 10:127 | 11:55 | SysteMHC | database |
| B4501 | 9:1427 | 10:377 | 11:169 | SysteMHC | database |
| B4901 | 8:34 | 9:301 | 10:33 | SysteMHC | database |
| B5001 | 9:219 | 10:51 | 11:17 | SysteMHC | database |
| B5201 | 8:87 | 9:52 | 10:5 | SysteMHC | database |
| B5501 | 8:36 | 9:254 | 10:59 | SysteMHC | database |
| B5601 | 9:195 | 10:80 | 11:41 | SysteMHC | database |
| B7301 | 9:103 | 10:53 | 11:44 | SysteMHC | database |
| C0301 | 8:10 | 9:78 | 10:14 | SysteMHC | database |
| C1204 | 8:11 | 9:66 | 10:8 | SysteMHC | database |
| C1505 | 8:6 | 9:21 | 10:3 | SysteMHC | database |
| A3301 | 8:780 | 9:288 | 10:37 | Bulik-Sullivan2018 | deconvolution |
| B1302 | 8:50 | 9:1325 | 10:156 | Bulik-Sullivan2018 | deconvolution |
| B1503 | 8:39 | 9:698 | 10:77 | Bulik-Sullivan2018 | deconvolution |

Table S2 Performance of cysteine frequency adjusted PSSMs on prediction of cysteine-free HLA-ligands.

| HLA alleles | 0.1%PPV on cysteine-free HLA-ligands | | |
|---|---|---|---|
| | Normal PSSM | Modified PSSM_1 | Modified PSSM_2 |
| A1101 | 0.352 | 0.346 | 0.327 |
| A0207 | 0.397 | 0.359 | 0.369 |
| A0201 | 0.348 | 0.33 | 0.344 |
| A2402 | 0.452 | 0.402 | 0.412 |
| A0203 | 0.436 | 0.431 | 0.438 |
| A0101 | 0.756 | 0.624 | 0.613 |
| A0301 | 0.449 | 0.439 | 0.433 |
| A3101 | 0.5 | 0.427 | 0.416 |
| B3501 | 0.445 | 0.386 | 0.368 |
| B4001 | 0.346 | 0.37 | 0.333 |
| B4403 | 0.581 | 0.526 | 0.513 |
| B5101 | 0.494 | 0.388 | 0.382 |
| B5401 | 0.492 | 0.358 | 0.35 |
| B0702 | 0.261 | 0.244 | 0.216 |
| A3201 | 0.566 | 0.557 | 0.494 |
| B1501 | 0.375 | 0.304 | 0.232 |
| A6802 | 0.475 | 0.437 | 0.416 |
| A0204 | 0.429 | 0.413 | 0.437 |
| A2902 | 0.588 | 0.475 | 0.441 |
| B4402 | 0.639 | 0.535 | 0.488 |
| B5701 | 0.454 | 0.31 | 0.29 |
| Average | 0.468 | 0.412 | 0.396 |
